# Fluorescence Microscopy of Piezo1 in Droplet Hydrogel Bilayers

**DOI:** 10.1101/508176

**Authors:** Oskar B. Jaggers, Pietro Ridone, Boris Martinac, Matthew A. B. Baker

## Abstract

Mechanosensitive ion channels are membrane gated pores which are activated by mechanical stimuli. The focus of this study is on Piezo1, a newly discovered, large, mammalian, mechanosensitive ion channel, which has been linked to diseases such as dehydrated hereditary stomatocytosis (*Xerocytosis*) and lymphatic dysplasia. Here we utilize an established *in-vitro* artificial bilayer system to interrogate single Piezo1 channel activity. The droplet-hydrogel bilayer (DHB) system uniquely allows the simultaneous recording of electrical activity and fluorescence imaging of labelled protein. We successfully reconstituted fluorescently labelled Piezo1 ion channels in DHBs and verified activity using electrophysiology in the same system. We demonstrate successful insertion and activation of hPiezo1-GFP in bilayers of varying composition. Furthermore, we compare the Piezo1 bilayer reconstitution with measurements of insertion and activation of KcsA channels to reproduce the channel conductances reported in the literature. Together, our results showcase the use of DHBs for future experiments allowing simultaneous measurements of ion channel gating while visualising the channel proteins using fluorescence.

## Introduction

The Piezo family are newly discovered mechanosensitive channels found in a wide range of mammalian cells and tissues. The family consists of both Piezo1 and Piezo2 channels found mainly in endothelial, urothelial, and renal epithelial cells [1]. The channel has been found to play a crucial role in the circulatory system due to its role in the sensing of shear-stress within blood vessel endothelial cells [2], and key roles in blood vessel formation, vascular tone regulation, red blood cell homeostasis, and the remodeling of small resistant arteries upon hypertension [3]. Due to the channel’s role in these key processes, mutations to the protein have been linked to several genetic disorders. Some examples include *Xerocytosis* (dehydrated hereditary stomatocytosis), where red blood cells are dehydrated [4], as well as lymphatic dysplasia, which causes persistent swelling due to lymphatic blockage [5]. A further point of interest is that the Piezo1 channel is extremely large in comparison to other mechanosensitive ion channels. While the bacterial mechanosensitive protein MscL is ~70 kDa in size [6], the Piezo1 protein is ~900 kDa [7].

The channel can be activated by a number of mechanical stimuli *in vitro*, such as cell stretching [8], as well as by using microfluidic chambers which can mimic sheer stress [9]. It is also ligand-gated. *Yoda1*, for example, is a known agonist which is lowering Piezo1’s mechanical sensitivity, thus allowing for easier activation [10]. Another molecule, *Dooku1*, has been found to antagonize *Yoda1*’s action [11]. The structure of Piezo1 was recently solved by multiple separate research groups at high resolution [6, 12-14] (Fig. 1).

**FIGURE 1.**
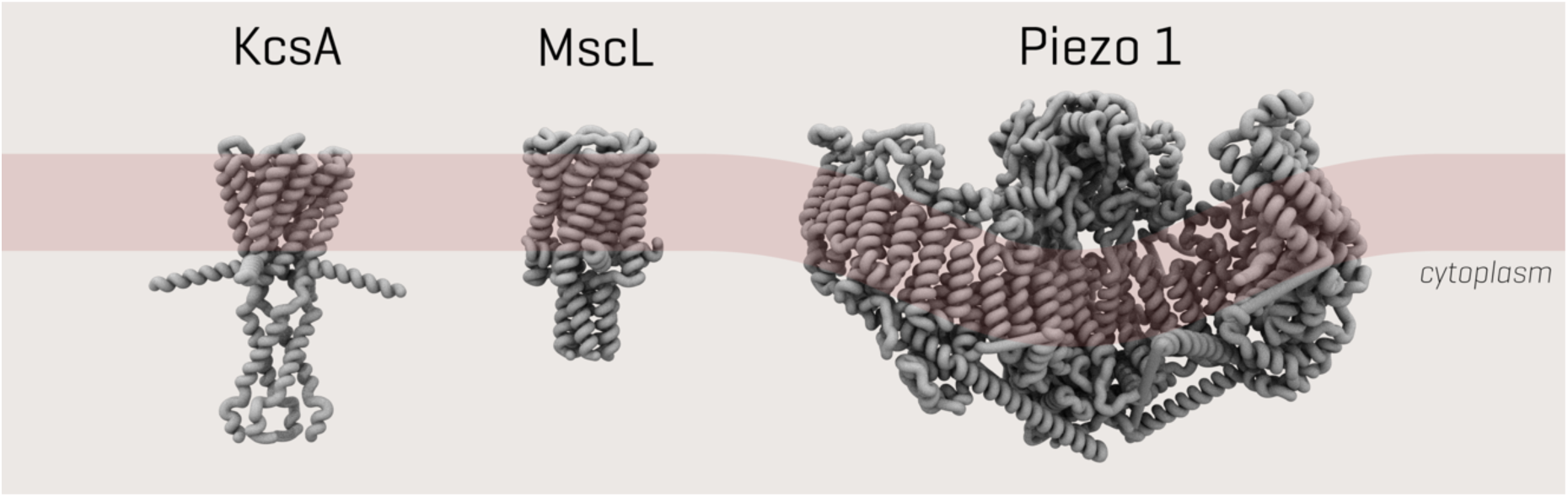
3D structure of ion channels in this study. Size and structure of ion channels KcsA, bacterial mechanosensory MscL and Piezo1 respectively. The coils indicate the alpha helices.

Artificial mimics and models are used to replicate membrane processes in a controllable manner. These *in vitro* methods allow for the analysis of the specific roles for each lipid in a complex lipid composition in terms of both function and transduction of force. Within the last 10 years, the latest approaches have moved from an aqueous environment towards aqueous droplets within an organic phase [15]. These droplet-based approaches have several advantages over previous methods, such as: ease of control of lipid asymmetry, the ability to optically image the bilayer, and the ease with which a wide range of proteins can be reconstituted [16]. We focus here on droplet hydrogel bilayers (DHBs), which are formed when an aqueous lipid droplet interfaces with a hydrogel, after both droplet and hydrogel have been incubated in lipids dissolved in hexadecane. This system allows for total control of lipid composition and asymmetry. Furthermore, this technique produces horizontally orientated bilayers, enabling simultaneous electrophysiology and fluorescent microscopy of ion channels reconstituted into the bilayers.

We performed here the first fluorescence microscopy of hPiezo1-GFP in DHBs. First, the background fluorescence of empty liposomes was measured. This allowed us to image hPiezo1-GFP with confidence that what we observed was representative of the protein, in conjunction with separate characterisation of the electrophysiology of the protein in our *in vitro* system. Successful integration of the channel led to further experiments investigating the effect of cholesterol on hPiezo1-GFP. We measured the electrical activity of lipid-only DHBs, KcsA in DHBs and hPiezo1-GFP in DHBs to showcase DHBs as a platform for combining imaging and activation of mechanosensitive ion channels.

## Results

### Fluorescence Microscopy of hPiezo1-GFP in DHBs

To generate droplet hydrogel bilayers, agarose at low concentration (0.75% w/v) was spin-coated onto a plasma-cleaned coverslip and then a fabricated PDMS device (Fig. 2A) was adhered to the coverslip using more agarose (3.2% w/v). Each of the 16 wells was filled with lipid in hexadecane (9.5 mg/mL) in order to assemble a lipid monolayer on the hydrogel surface. Separately, an aqueous droplet was incubated in an identical solution of lipid hexadecane to form a lipid monolayer around the droplet, and then this droplet was brought into contact with the monolayer on the hydrogel to form a bilayer (Fig. 2BC), and an electrode was inserted for electrophysiological recording. If the droplet contained proteoliposomes, i.e., azolectin liposomes containing reconstituted channels (Fig. 2D), these then fused with the bilayer (Fig. 2E) and the membrane protein was inserted where it could be interrogated by fluorescence and electrical recording (Fig. 2F). For a full description, see Materials and Methods. To characterize the background fluorescence, we first generated droplet hydrogel where the interior of the droplet was a suspension of azolectin-only liposomes (Fig. 2). Upon illumination with 488 nm laser, there was a low level of diffuse autofluorescence, but no distinct puncta from diffusing single molecule fluorophores (Fig. 3A).

**FIGURE 2.**
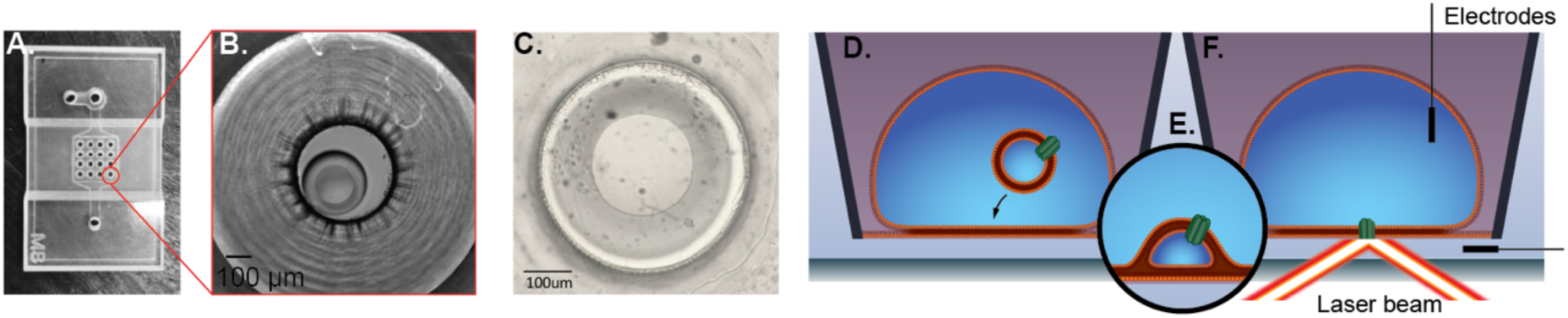
Droplet Hydrogel Bilayers (DHBs). PDMS device (A) with 16 wells is adhered to an agarose-coated size-matched 22 mm x 40 mm coverslip and each well is filled with DPhPC/hexadecane mixture to form a lipid monolayer on the surface. Aqueous droplets consisting of a suspension of proteoliposomes (azolectin liposomes reconstituted with ion channel proteins) are separately incubated in a DPhPC/hexadecane mixture to form a monolayer. The droplet monolayer is then joined with a monolayer on an agarose hydrogel on a glass coverslip to form a bilayer oriented horizontally on a transparent surface (bright field micrograph at low magnification: B; and high magnification: C). Proteoliposomes inside the droplet (D) fuse spontaneously with the bilayer (E) to insert membrane protein, and then an electrode can be inserted in the droplet to record electrical activity whilst simultaneously imaging the bilayer with fluorescence microscopy (F).

**FIGURE 3.**
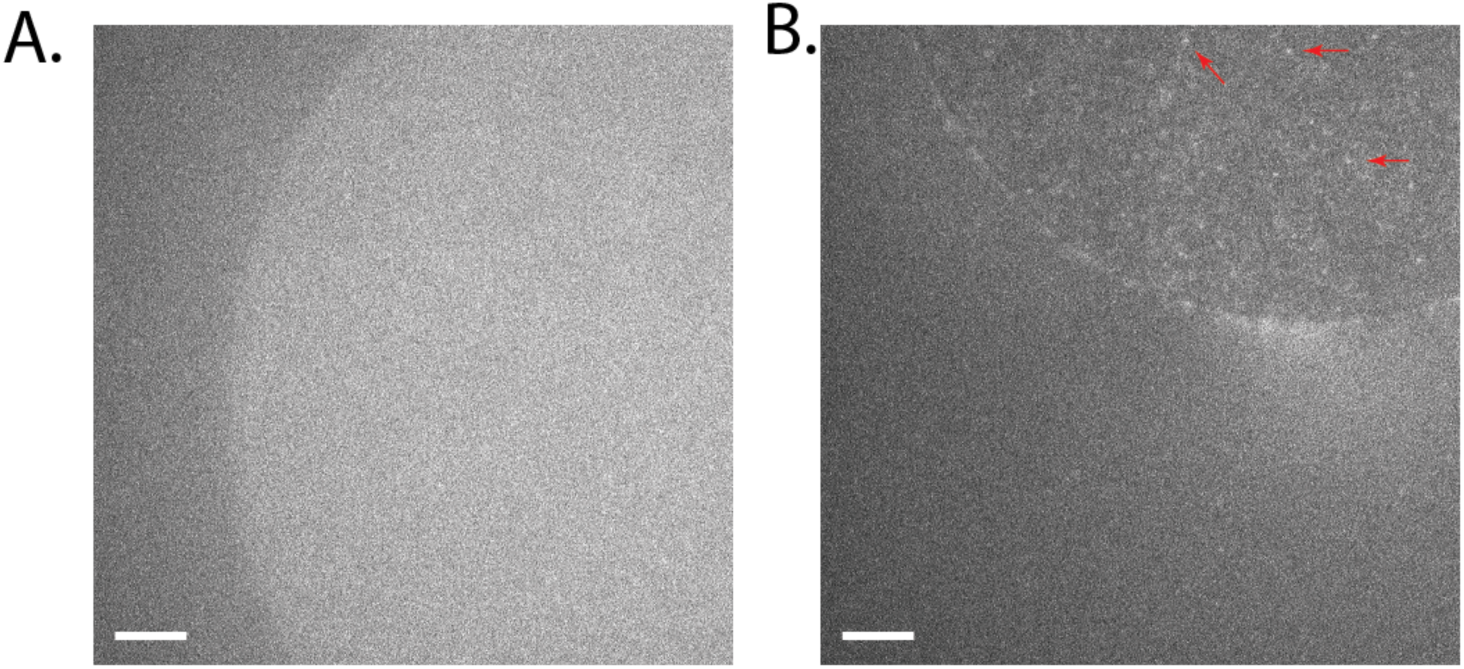
Fluorescence of hPiezo1-GFP in DHBs. (A) Autofluorescence from a DPhPC bilayer with interior consisting of empty azolectin liposomes. (B) Fluorescence of hPiezo1-GFP inserted in bilayer diffusing throughout the bilayer. Example puncta indicated with red arrows. Diffusion can be seen more clearly in Supplementary Movies 1 & 2. Exposure time: 100 ms; Laser power: 0.2 µW/um^2^; Image brightness scaled to range from 300-1600 a.u, on Andor EMCCD Camera (see Methods). Scale bar indicates 10 um.

### Fluorescence of hPiezo1-GFP in DHBs

Following background controls, hPiezo1-GFP was imaged under the same conditions (see Methods). Individual fluorescent puncta were observed (Fig. 3), especially in the central region of the DHB, corresponding to the fully formed dual-leaflet bilayer. In a frame-by-frame video the contrast in movement and diffusion in the plane of the bilayer of hPiezo1-GFP was clearly observable (Supplementary Movies 1 & 2).

### Effect of Cholesterol hPiezo1-GFP in DHBs

Cholesterol has been shown to influence the clustering of Piezo1[17]. Thus, we sought to determine the effect of cholesterol on Piezo1 in DHBs. We imaged the hPiezo1-GFP that was prepared from azolectin liposomes into DHBs composed of 8:92 cholesterol:DPhPC. The fluorescence of hPiezo1-GFP observed in (Figure 3B) has tenfold fewer puncta per square micrometre than in the absence of cholesterol (Figure 4A). Diffusion and movement can be observed in Supplmentary Movies 3 & 4.

**FIGURE 4:**
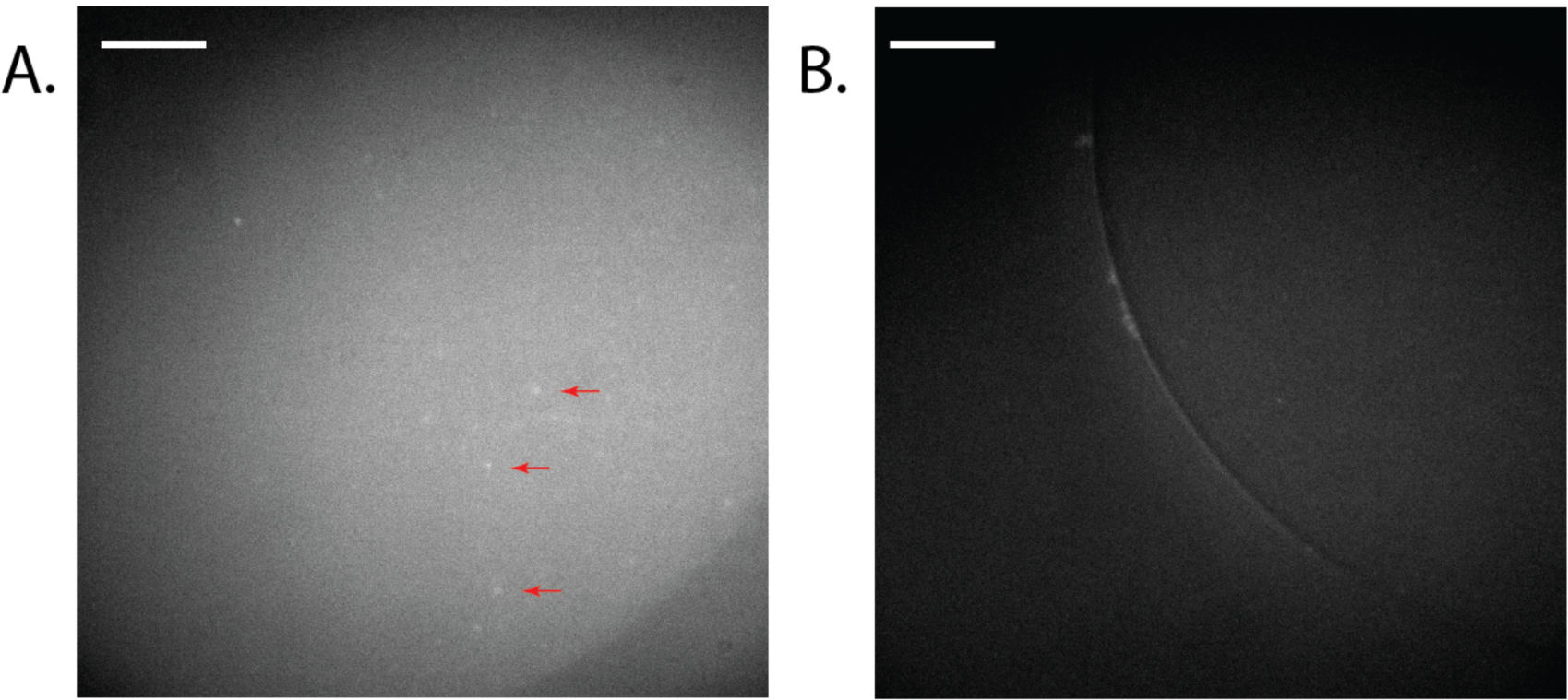
Fluorescence of hP1-GFP in DHBs with 8% cholesterol. (A) hP1-GFP diffusing in a bilayer composed of 8:92 cholesterol to DPhPC (see Methods). Example puncta are indicated by red arrows. (B) Positive control for fluorescence, but not membrane protein insertion, observed using a DPhPC bilayer with an interior of GFP-filled azolectin liposomes (fluorophore present, but no membrane protein). Background fluorescence is observable but not individual puncta diffusing inside the bilayer. Diffusion can be seen more clearly in Supplementary Movies 3 & 4. Exposure time: 100 ms; Laser power: 0.3 µW/um^2^; Image brightness scaled to range from 150-300 a.u on Hamamatsu 95B camera. Scale bar indicates 20 µm.

### Insertion and Activation of ion channels in DPhPC bilayers

#### KcsA as a pH-sensitive positive control

KcsA, a pH-sensitive bacterial channel was reconstituted in azolectin liposomes at a ratio of 1:250 protein to lipid as a positive control. Acidic conditions (200 mM KCL, 5 mM HEPES, pH 4.2) triggered activity (Fig. 5) with two current levels indicating the channel activity. The first gating level was at 4 pA, with the second level at 8 pA corresponding to a single channel conductance of ~20 pS[18].

**FIGURE 5:**
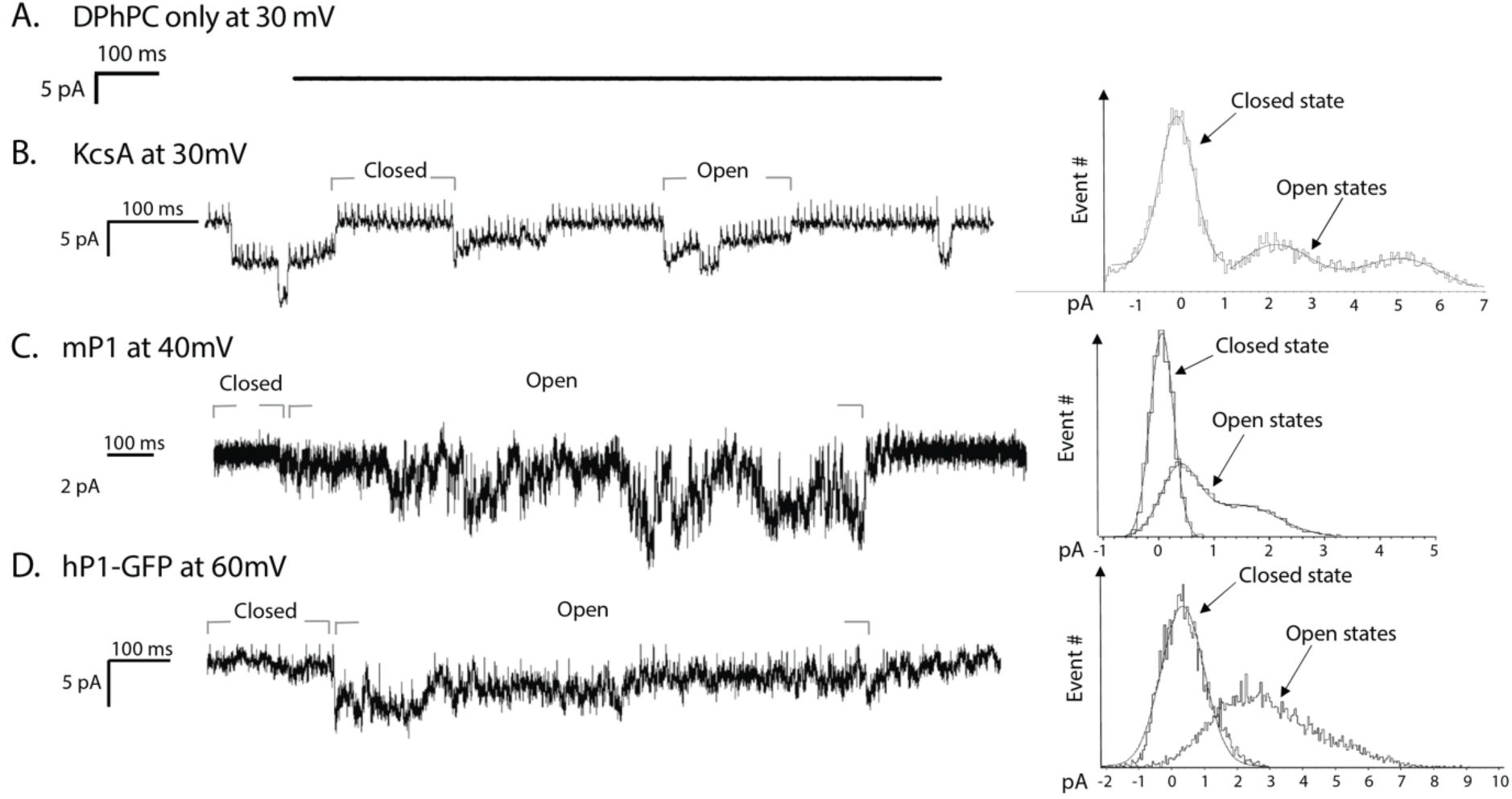
Single channel recordings of Piezo1 in DHBs. (A) Empty liposomes under the application of 30 mV do not show channel activation. (B) KcsA at pH 4.0 and application of 30 mV shows closed and open states. (C) mPiezo1 under application of 40 mV. (D) hPiezo1-GFP upon application of 60 mV. Histograms of open and closed states defined by labelled grey bars are shown for each protein on the right.

#### hPiezo1 activity in recorded from liposomes and DHBs

The human Piezo1 (hPiezo1), was reconstituted into Azolectin liposomes at the 1:250 protein to lipid ratio and incorporated in DPhPC bilayers. When 30 mV was applied to the system, gating like activity was observed. This activity was sporadic and had several levels from 3 pA up to a maximum of 18 pA, corresponding to a single channel conductance of ~100 pS.

For comparison, the hPiezo1-GFP currents were recorded from the channels reconstituted into azolectin liposomes using patch clamp technique (see Methods). During the application of 40 mV to the patched liposomes, there were several single channel events showing multiple current levels (Fig. 6) corresponding to a single channel conductance of ~100 pS, similar to the current levels recorded using the DHB system. In addition, apparently two channels were gating cooperatively in the shown trace.

**FIGURE 6.**
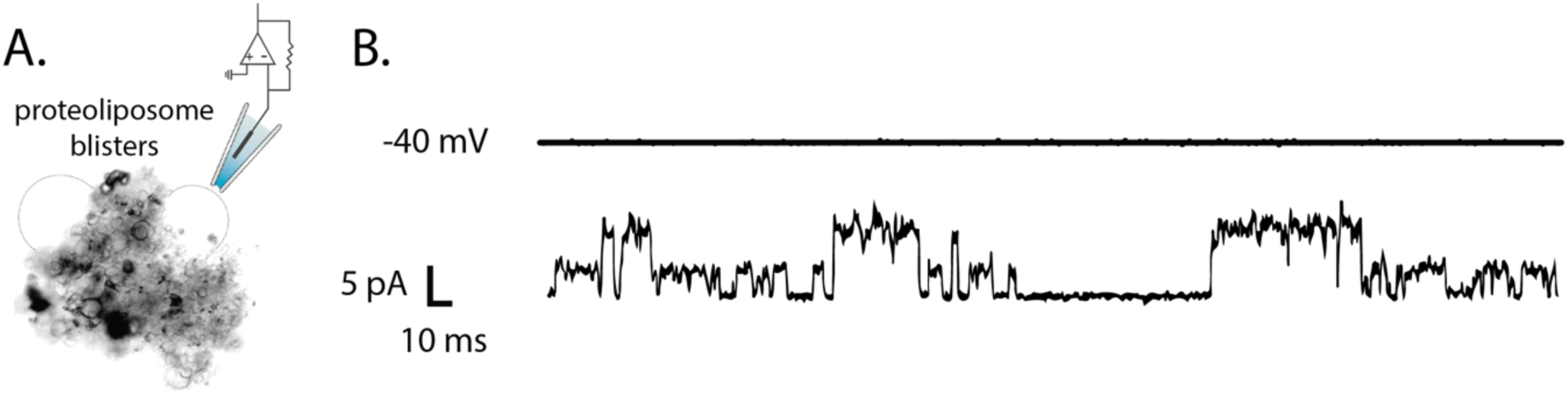
Spontaneous activation of human Piezo1 reconstituuted into liposomes. (A) Azolectin proteoliposomes containing hPiezo1-GFP were patched using patch-clamp apparatus where voltage and stretch can be applied simultaneously. B) During application of −40 mV, activity of hPiezo1 was observed.

## Discussion

We successfully integrated hPiezo1-GFP into the membrane of artificial bilayers, verifying this with fluorescence and verifying activity using electrophysiology. One of the key purposes behind using the DHB system is that the bilayers are horizontally oriented and thus can be interrogated using simultaneous electrophysiology and fluorescent microscopy. Here we have done each measurement in isolation. The next step will be to combine these measurements and image the protein while recording activity to correlate electrical activity with spatial location, stoichiometry, and, in dual-label samples, to correlate conformational change using single molecule Förster Resonance Energy Transfer (smFRET)[19].

We observed single molecule diffusion in and out of the membrane, and corroborated cell measurements that have co-localised cholesterol with Piezo1. This needs to be examined in further detail as lipid composition becomes adjusted to include other natural mammalian lipids such as phosphatidylserine and phosphatidylinositol, which are present in the inner leaflet of the bilayers of cell membranes. In conjunction with other work, a picture is forming that suggests cholesterol is essential for efficient mechanotransduction by Piezo1. Depletion of cholesterol affects channel activation and latency[20]. Being able to simultaneously perform electrophysiology and fluorescent microscopy with DHBs, would help to link this clustering to activity. We observed (Fig. 4) that the fluorescent spots in bilayers, corresponding to hPiezo1-GFP complexes, were larger but fewer in number in the presence of cholesterol. This supports the elsewhere observed clustering of Piezo1 in cholesterol-dependent domains[21].

We used KcsA as a positive control for electrical activity as also previously reported[18]. We verified that at pH 8 there was no observed activity and recorded gating events at pH 4.0 (Fig. 5), with single channel currents ranging from 4-8 pA. This value is comparable with the patched liposomes containing KcsA from the literature of around 100 picoSiemens (pS) [22]. Subsequently, hPiezo1 protein was also successfully inserted and activated in DHBs. We observed single channel currents ranging from 3-6 pA (Fig. 4). With an applied voltage of 30 mV this equates to a single channel conductance of around 100 pS. There is no established conductance of the protein to use for comparison currently. However, our measurements do correlate well with our own recordings of hPiezo1-GFP via patch clamping of liposomes (Fig. 5), where we measured a conductance of ~100 pS as well.

## MATERIALS AND METHODS

### Protein Purification

HEK293T cells were cultured in 150 cm^2^ flasks until ~70% confluent in Dulbecco’s Modified Eagle’s Medium (DMEM) supplemented with 10% Newborn Calf Serum (SIGMA) and then transfected with pCGFP-EU hP1 6his-Flag plasmid (40 ug per flask, each containing 20 ml of culture media) encoding the human WT Piezo1 channel fused at the N-terminus to GFP and followed by a 6xHistidine tag and a FLAG tag for affinity chromatography. Transfection was performed using 160 ug of Polyethyleneimine (PEI), incubated with plasmid DNA at room temperature for 30 min in 1 mL OptiMEM media, then added to each culture flask. After 48 h, the transfected cells were collected, washed twice with PBS and homogenized in buffer A, containing 25 mM Tris-HCl pH 7.2, 140 mM NaCl, 2 mM TCEP and a cocktail of protease inhibitors (Roche) using a syringe fitted with a 26G ½ needle. The resulting homogenate was then adjusted with detergents CHAPS (1%), C12E9 (0.1%) and 0.5% (w/v) L-a-phosphatidylcholine (Avanti) and incubated at 4°C for 3 h with gentle shaking. After centrifugation (12000 rpm, 20 min, 4°C), the supernatant was collected and incubated with anti-FLAG M2 affinity Gel (A2220, SIGMA) at 4°C overnight. The resin was washed extensively with buffer B, 50 mM Tris-HCl pH 7.2, 150mM NaCl, 2 mM TCEP, protease inhibitors, 0.6% CHAPS, 0.025% C12E9 and 0.14% (w/v) L-a-phosphatidylcholine. Bound hPiezo1-GFP was eluted overnight with buffer B plus 3xFLAG peptide (0.1 mg/ml). Evaluation of sample quality was carried out using poly-acrylamide gel electrophoresis (BIORAD) and fluorescent size-exclusion chromatography (FSEC) (Superpose-6 10/300 GL, GE Healthcare) using buffer C (50mM Na-PIPES, pH 7.2, 150 mM NaCl, 2 mM TCEP, 0.3% CHAPS, 0.025% C12E9 and 0.07% (w/v) L-a-phosphatidylcholine). Mouse Piezo1 (mPiezo1) was purified as previously in [23]

### Proteoliposome Preparation

To reconstitute ion channels into the droplet-hydrogel system, proteoliposomes were prepared prior to droplet incubation. Liposomes were created using a dehydration/rehydration method [18], followed by incubation with detergent solubilized protein and biobeads.

### Liposome formation

Lipid (2 mg) was dissolved in chloroform in a glass vial. The chloroform was evaporated under nitrogen flow, forming a layer of lipid film. The experimental buffer (1 mL, 200 mM KCl, 5 mM HEPES, pH 7.4) was added to the vial, followed by vortexing and sonication (15 min). The solution was then passed through a lipid vesicle extruder (AVESTIN, 100 nm).

Liposome samples prepared: 100% Soybean Azolectin (P5638, Sigma), 100% DPhPC (Avanti, 850356), Azolectin: Cholesterol (80:20; w:w; Cholesterol C8667, Sigma)

For samples not containing DPhPC, the experimental buffer (1 mL, 200 mM KCl, 5 mM HEPES, pH 7.4) was added to the vial, followed by vortexing and sonication (15 min). Conversely, samples containing DPhPC were processed in the same way using a 0.5mM KCl experimental buffer instead.

### Protein reconstitution

The protein of choice (KcsA, mPiezo1 or hPiezo1) was added in a 1:250 protein: lipid ratio. This was incubated on a rotary table (1 h, 4°C), followed by the addition of biobeads. The biobeads were exchanged three times over a 19 h period (after incubation periods of 2 h, 2 h, 15 h). After biobead removal, the proteoliposome solution was diluted (10x or 100x) in the experimental buffer (1 mL, 200 mM KCl, 5 mM HEPES, pH 7.4). This final solution was used for the droplet formation.

### Hydrogel preparation

Deionized water (1 mL) was added to Agarose (7.5 mg), producing a 0.75% (w/v) solution. The desired experimental buffer (1 mL) was added to Agarose (32.5 mg), producing a 3.25% (w/v) solution. Both agarose solutions were used for device assembly as described later.

### Lipid preparation

Lipid in hexadecane (9.5 mg/mL) is used to flow through the device, forming a lipid monolayer at the base of each well. In order to produce this solution of DPhPC in Hexadecane with a final concentration of (9.5 mg/mL), 190μL of stock DPhPC in chloroform (50 mg/mL) was added to a glass vial. The chloroform was removed forming a lipid film by nitrogen flow and vacuum desiccation (30 min). Hexadecane (1 mL) was added to the vial, followed by vortexing and sonication (15 min). For DPhPC/cholesterol experiments, cholesterol (C8667, Sigma) was prepared as 100 mM stock in chloroform and mixed at 8% with DPhPC to form final composition of 8:92 cholesterol:DPhPC.

### Device Assembly

The DHB device was made of PMMA and contains sixteen 1 mm wide wells with a microfluidic matrix surrounding the wells from underneath. A glass coverslip was spin coated (Laurell Technologies Corporation® 4000rpm, 30s) with agarose (0.7% (w/v), 120 uL, 90°C), creating a thin film on the surface. The coverslip was pressed on the underside of the device, followed by the addition of agarose (3.25% (w/v), 140 uL, 90°C) flowed from above, sealing the coverslip. The wells were then filled with 60 uL of the previously prepared lipid in hexadecane.

### Droplet Incubation

Meanwhile, the DHB incubation chamber (PMMA, 10 grooves) was filled with lipid in hexadecane (9.5 mg/ml). The proteoliposome solution prepared earlier were transferred with a pipette (0.15 uL) into the grooves to be incubated, forming a lipid monolayer (20 min).

### Droplet Hydrogel Bilayer Formation

Once both lipid monolayers have formed, on the surface of the hydrogel at the bottom of the wells and around the resuspended droplets, the droplets are transferred into the wells. Upon contact the lipid monolayers will join together, forming a bilayer.

### Electrophysiology

The electrode was prepared by immersing silver wire in sodium hypochlorite (4%) forming an AgCl coating. The ground electrode was inserted into the agarose (3.25% (w/v) while the electrode was coated in a thin layer of agarose before insertion into a droplet with a bilayer. The electrodes were connected to a Bilayer Clamp Amplifier (BC-535, Warner Instruments), filtering the current at 1 kHz and receiving at 5 kHz with a digitizer acquisition system (Digidata 1440 A) and acquisition software (pCLAMP10 by Molecular Devices, Sunnyvale, CA).

### Patch-Clamp Electrophysiology

The patch clamp experiment was carried out and analyzed using standard procedures as previously described[24, 25]. Briefly, dried proteoliposome samples were rehydrated using 200mM KCl experimental buffer and incubated at 4°C for a minimum of 3 hrs. 5 µl of rehydrated proteoliposomes were transferred to a patch-clamp recording chamber filled with 1 mL of patch clamp buffer (200 mM KCl, 5 mM HEPES, 40 mM MgCl_2_, pH =7.4). Recordings from liposomes were performed using the same patch buffer in the glass pipette and the bath. The channel currents were recorded with an AxoPatch 200B amplifier (Axon Instruments) in the inside-out patch configuration, and data were collected at 10 kHz sampling rate with 2 kHz filtration. Borosilicate glass pipettes (Drummond Scientific, Broomall, PA) with bubble number 5.0-6.0 (equivalent to 2-4 MΩ resistance in the patch clamp buffer) were pulled using a Narishige puller (PP-83; Narishige). Data collection and analysis were carried out using the pClamp10 software suite (Axon Instruments).

### Imaging

Fluorescence micrographs in Fig. 3 were imaged using TIRF illumination on a Nikon Ellipse I95. Laser excitation was supplied by Vortran Stradus laser diode units (488 nm), focused onto the back focal plane of a Nikon 100x Plan Fluor objective, and images were recorded on a Photometrics Prime 95B Scientific CMOS camera. For fluorescence micrographs in Fig. 4, the same laser and objective were used, however the images were recorded on an Andor EMCCD 888 camera with gain set to 300. The exposure time for all images was 100 ms, with a laser flux of 0.2-0.3 µW/um^2^ (as noted in captions).

## Author Contributions

OBJ collected and analysed data and wrote the manuscript. PR collected and analysed data and wrote the manuscript. BM supervised the project and wrote the manuscript. MABB conceived and supervised the project, collected and analysed data, and wrote the manuscript. We acknowledge the assistance of Dr Yury Nikolaev and Mr Jon Berengut with preparation of Figure 1 and Figure 2 respectively.

