## Supplementary Movies - Captions for "Fluorescence Microscopy of Piezo1 in Droplet Hydrogel Bilayers"

**Supplementary Movie 1. Background autofluorescence in DHBs**. Autofluorescence from a DPhPC bilayer with interior consisting of empty azolectin liposomes. Exposure time: 100 ms; Laser power: 0.2 μW/um$^2$; Image brightness scaled to range from 300-1600 a.u, on Andor EMCCD Camera (see Methods). Whole image is 88.68 μm by 88.68 μm. Movie is played at 10 fps (100 ms exposures) with JPEG compression.

**Supplementary Movie 2. Fluorescence of hPiezo1-GFP inserted in DHB**. hPiezo1-GFP can be observed diffusing throughout the bilayer. Exposure time: 100 ms; Laser power: 0.2 μW/um$^2$; Image brightness scaled to range from 300-1600 a.u, on Andor EMCCD Camera (see Methods). Whole image is 88.68 μm by 88.68 μm. Movie is played at 10 fps (100 ms exposures) with JPEG compression.

**Supplementary Movie 3. Fluorescence of hP1-GFP in DHBs with 8% cholesterol**. hP1-GFP diffusing in a bilayer composed of 8:92 cholesterol to DPhPC. Exposure time: 100 ms; Laser power: 0.3 μW/um$^2$; Image brightness scaled to range from 150-300 a.u on Hamamatsu 95B camera (see Methods). Whole image is 132 μm by 132 μm. Movie is played at 10 fps (100 ms exposures) with JPEG compression.

**Supplementary Movie 4. Fluorescence of GFP in solution above bilayer of DHBs**. Positive control for fluorescence, but not membrane protein insertion, observed using a DPhPC bilayer with an interior of GFP-filled azolectin liposomes (fluorophore present, but no membrane protein). Exposure time: 100 ms; Laser power: 0.3 μW/um$^2$; Image brightness scaled to range from 150-300 a.u on Hamamatsu 95B camera (see Methods). Whole image is 132 μm by 132 μm. Movie is played at 10 fps (100 ms exposures) with JPEG compression.
